# Proficiency-Dependent and Dynamic Reorganization of Brain Networks During Children’s Second Language Learning

**DOI:** 10.64898/2025.12.10.693259

**Authors:** Nicole H. Skieresz, Sandy C. Marca, Micah M. Murray, Nicolas Rothen, Thomas P. Reber

**Author notes:** Correspondence concerning this article should be addressed to Nicole H. Skieresz. These authors contributed equally to this work (Micah M. Murray, Nicolas Rothen, Thomas P. Reber).

## Abstract

Children’s second language (L2) vocabulary learning is often indexed with event-related potentials (ERPs) by the N400 incongruity effect, yet its stability in real classrooms is unclear. We recorded 64-channel EEG from 42 children (aged 9-15 years) during a translation-recognition task before and after a regular instructional vocabulary unit and sub-grouped children by baseline proficiency. We quantified the canonical N400 (300–500 ms, centroparietal region of interest; ROI) alongside electrical-neuroimaging indices of ERP strength and topography (Global Field Power, Global Map Dissimilarity, ERP template-map preponderance), using adult template maps from a prior study as a normative reference. Behaviorally, children showed large gains in vocabulary accuracy and higher discrimination sensitivity (*d′*) in post-assessment. The classical N400 incongruity effect did not increase from pre- to post-learning. By contrast, global and topographic measures were learning–sensitive: Global Field Power increased across sessions and interacted with proficiency, and Global Map Dissimilarity showed main effects of session and proficiency as well as their interaction, indicating proficiency-dependent topographic reorganization. Adult normative template maps backfitted well; within the N400 window, high-proficiency children more consistently occupied an adult-like configuration; later (∼650–800 ms), patterns also differentiated by proficiency and congruency. Relative to adults, children exhibited more diffuse topographies and stronger dependence on individual proficiency. In classroom settings, children’s L2 vocabulary learning is therefore more sensitively indexed by scalp topography and global field dynamics than by a single N400 amplitude, positioning electrical neuroimaging as a proficiency-dependent, classroom-relevant marker of L2 acquisition.

## 1 Introduction

Children’s ability to acquire new vocabulary in a second language (L2) undergoes substantial developmental changes, supported by the maturation of neural substrates underlying semantic processing and cognitive control (Friederici, 2006; Neville et al., 1993; Schlaggar et al., 2002). Electrophysiological studies in adult learners have consistently identified robust neural markers associated with successful semantic integration, prominently including the N400 event-related potential (ERP; Kutas & Federmeier, 2011; Van Petten & Luka, 2012). The N400 is a negative-going ERP component peaking around 400 ms after word onset, which is sensitive to semantic processing and typically increases (i.e., becomes more negative) for words that are incongruent or unexpected within a given context—a phenomenon known as the N400 incongruity effect, commonly studied in second language research using translation recognition tasks. In such tasks, participants respond to word pairs that are either correct (congruent) or incorrect (incongruent) translations, with larger N400 amplitudes for incongruent pairs reflecting increased semantic integration demands (cf. Midgley et al., 2009; Pu et al., 2016). Additionally, global EEG features such as ERP template-map analyses (topographic clustering), Global Field Power (GFP), and Global Map Dissimilarity (GMD) have provided insights into the spatiotemporal dynamics of semantic processing in adults (Koenig et al., 2011; Michel & Koenig, 2018; Michel et al., 2004). However, whether these neural markers reliably generalize to children in natural educational settings remains unclear, particularly with regard to developmental and individual variability in authentic classroom contexts.

Research on the N400 component in childhood has documented several developmental variations, including delayed peak latencies, altered amplitudes, and less focal scalp distributions relative to adults (Holcomb et al., 1992; Juottonen et al., 1996; Friedrich & Friederici, 2004; Hahne et al., 2004; Skeide et al., 2015; Benau et al., 2011). For example, Holcomb et al. (1992) showed that children aged 7 to 12 years exhibit a later N400 peak—often by ∼100 ms—alongside broader scalp distributions. Juottonen et al. (1996) found that while children’s N400s are more anterior and diffuse, the incongruity effect was significantly larger than in adults. Skeide et al. (2015) similarly report broader and often larger N400 effects during early school age. Benau et al. (2011) documented that 7–10-year-olds show a larger and more widespread N400 effect compared to adults, with delayed peaks and anterior topography. Friedrich and Friederici (2004) confirmed a trajectory where N400 amplitude and timing become more adult-like with age – specifically showing increased sensitivity to semantic incongruency and a posterior shift in topography.

Importantly, a growing body of work highlights pronounced individual differences in N400 development among children, depending on language proficiency, vocabulary size, and task performance. Children with higher language competence or reading skills tend to show more focal, adult-like N400 responses, while those with lower proficiency often display larger, more diffuse, and prolonged N400 effects (Benau et al., 2011; Coch & Holcomb, 2003; Hahne et al., 2004; Skeide et al., 2015; Weber-Fox & Neville, 1996). Such findings indicate that the neural mechanisms of semantic processing in childhood are shaped not only by age, but also by individual differences in language and cognitive development.

These developmental and individual differences in the N400 are thought to reflect the progressive refinement and increased efficiency of neural networks involved in semantic access and integration. In particular, the initially broad and diffuse N400 responses observed in children likely result from less specialized or less efficiently interconnected language-related brain regions, whereas the gradual shift toward a more focal and posterior distribution with age and competence signals the maturation and functional specialization of left temporo-parietal areas central to adult-like semantic processing (cf. Friederici, 2006; Skeide et al., 2015).

In a recent ERP study conducted with adult learners, significant learning-related changes in the N400 incongruity effect were observed accompanied by systematic modulations of ERP template maps, Global Field Power (GFP), and Global Map Dissimilarity (GMD) following vocabulary acquisition (Skieresz et al., 2025). These findings demonstrate the sensitivity of electrical-neuroimaging markers to semantic integration processes in controlled laboratory conditions. The present study backfits adult normative ERP template maps from the previous study (Skieresz et al., 2025) to children’s EEG data, allowing for a systematic, quantitative comparison of ERP template-map preponderance—the proportion of epoch time frames assigned to a given template map—within defined processing epochs across age groups. By focusing on the temporal properties, this study provides a fine-grained assessment of developmental continuity or divergence in the stability and engagement of semantic processing-related brain states.

Classroom-based learning offers a unique and ecologically valid setting for neurocognitive research, presenting challenges and complexities not found in controlled laboratory environments (Ansari & Coch, 2006; Coch & Holcomb, 2003). In real-world educational contexts, learners contend with an interplay of cognitive, motivational, social, and attentional demands, as well as exposure to variable stimuli, peer interactions, and instructional strategies. These factors may impact not only the depth of semantic processing, but also the reliability and temporal dynamics of EEG markers. For example, everyday distractions, fluctuating engagement, and classroom heterogeneity can influence attention and resource allocation, thereby affecting electrophysiological indices such as the N400 and ERP template-map measures (alongside GFP and GMD). Additionally, more variable and authentic motivational and affective states in classroom settings may further shape neural responses in ways not fully captured by laboratory paradigms. Converging classroom EEG work demonstrates that robust neural signatures can be captured in situ and relate to engagement and later memory for instructional material (Dikker et al., 2017).

Consequently, assessing electrophysiological markers within naturalistic educational settings is crucial for establishing the ecological validity and practical relevance of neuroscience findings in real-world learning. Only by validating neural markers such as the N400 effect and ERP template-map analyses under authentic classroom conditions can we ensure that these measures genuinely reflect the neurocognitive processes relevant for language learning and academic development. Such work lays the foundation for translational educational neuroscience and informs the development of age-appropriate, evidence-based interventions and assessments.

Against this backdrop, we examined whether adult normative template maps defined for the N400 time window can be backfitted to children’s ERPs during classroom-based L2 vocabulary learning, permitting quantitative assessment of template-map preponderance. We further asked to what extent neurophysiological correlates of L2 vocabulary learning parallel concrete behavioral improvements across sessions, considering both a canonical centroparietal N400 amplitude and global/topographic indices (GFP, GMD, ERP template-map preponderance) as candidate neural measures of learning-related change.

## 2 Methods

### 2.1 Participants

Sixty-five children (34 females; *M*_age_ = 10.7, *SD*_age_ = 1.2, age range = 9–15 years) participated in this study as part of a larger classroom-based project conducted at public primary schools in the canton of Valais, Switzerland. All participants were enrolled in either 4th grade or 6th grade. Of these, 52 children were native German speakers learning French as a second language (L2), and 13 were native French speakers learning German as a second language (see *Table 1* for class-wise breakdown; see also *Supplementary Table S2A* for detailed characteristics). Parental written informed consent was obtained for all participants, and each child received a personalized certificate of participation. For EEG assessment, a subsample of 42 children (19 females; *M*_age_ = 10.6 years, *SD*_age_ = 1.2, age range = 9-15 years) completed both pre- and post-intervention measurements. The study was approved by the ethics committee of UniDistance Suisse (Nr. 2019-12-00002).

**Table 1.**
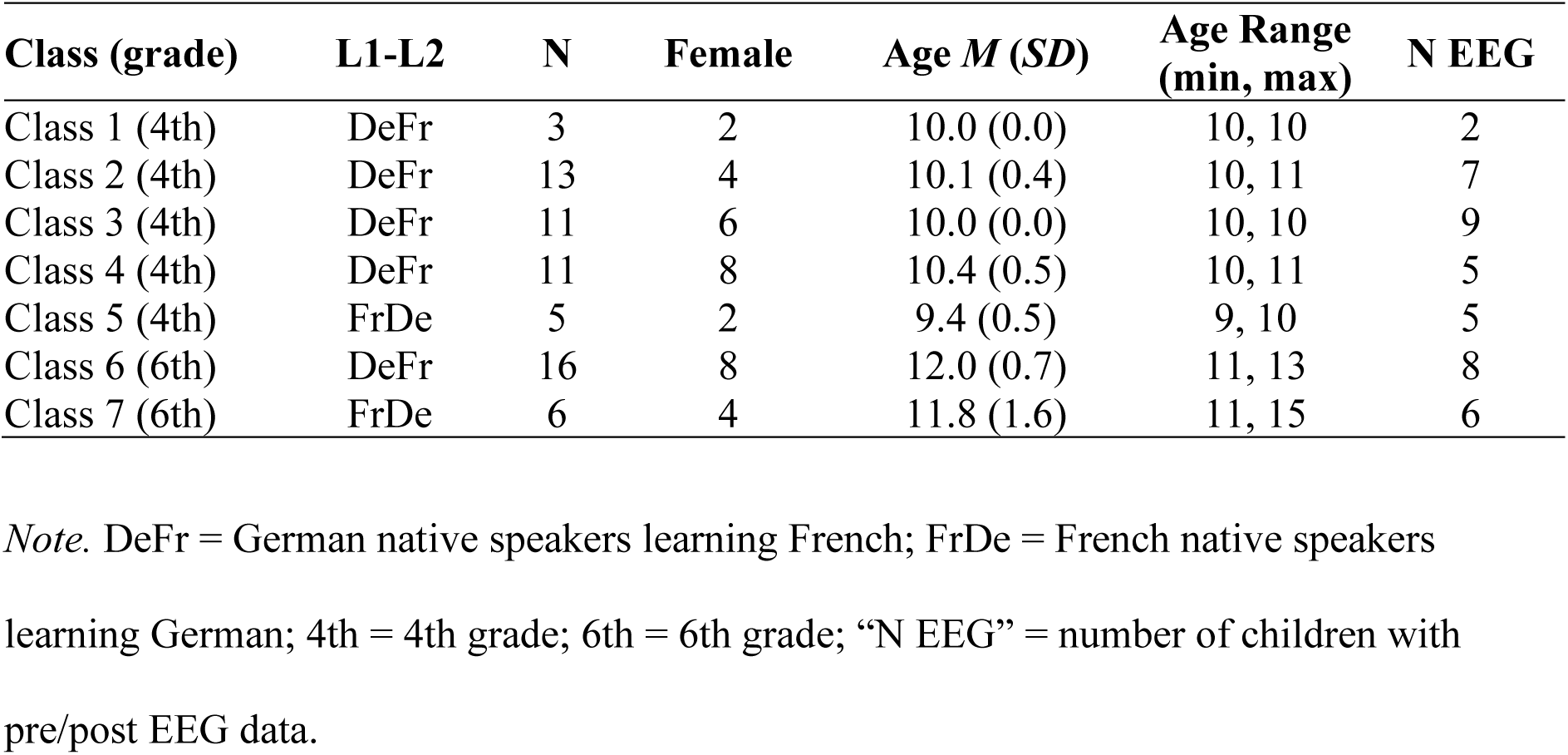
Demographic and sample characteristics by class.

| Class (grade) | L1-L2 | N | Female | Age $M$ ($SD$) | Age Range (min, max) | N EEG |
| --- | --- | --- | --- | --- | --- | --- |
| Class 1 (4th) | DeFr | 3 | 2 | 10.0 (0.0) | 10, 10 | 2 |
| Class 2 (4th) | DeFr | 13 | 4 | 10.1 (0.4) | 10, 11 | 7 |
| Class 3 (4th) | DeFr | 11 | 6 | 10.0 (0.0) | 10, 10 | 9 |
| Class 4 (4th) | DeFr | 11 | 8 | 10.4 (0.5) | 10, 11 | 5 |
| Class 5 (4th) | FrDe | 5 | 2 | 9.4 (0.5) | 9, 10 | 5 |
| Class 6 (6th) | DeFr | 16 | 8 | 12.0 (0.7) | 11, 13 | 8 |
| Class 7 (6th) | FrDe | 6 | 4 | 11.8 (1.6) | 11, 15 | 6 |

*Note.* DeFr = German native speakers learning French; FrDe = French native speakers learning German; 4th = 4th grade; 6th = 6th grade; “N EEG” = number of children with pre/post EEG data.

### 2.2 Design

The study employed a mixed factorial design with both within- and between-subjects factors, following the same methodological structure as our previous study in adults (Skieresz et al., 2025). Within-subjects factors included Session (pre- vs. post-learning phase) and Congruency (congruent vs. incongruent translation pairs). The between-subjects factor was Proficiency Group. Given that all children already had L2 knowledge from regular classroom instruction, proficiency was operationalized at baseline: within the EEG subsample, a median split of pre-intervention vocabulary-test accuracy classified participants into low- and high-proficiency groups (ties at the median were assigned to the high-proficiency group). The study comprised three sequential phases: pre-learning assessment, vocabulary learning phase, and post-learning reassessment. Dependent variables included vocabulary-test accuracy (percent correct) and response-bias measures from the translation recognition task (sensitivity, *d′*; criterion, *c*; likelihood ratio, ln *β*). Neurophysiological measures included the mean N400 amplitude (300–500 ms, six channels), ERP template-map preponderance (proportion of time frames assigned ERP template maps based on spatial correlation within predefined epochs), Global Field Power (GFP), and Global Map Dissimilarity (GMD). GFP and GMD were computed frame-wise (millisecond-by-millisecond) across the 0-796 ms epoch.

### 2.3 Materials

#### 2.3.1 Stimuli

A randomly selected subset of 20 vocabulary items from the regular classroom curriculum, as planned and provided by the respective teachers for the instructional unit, was used for both the vocabulary test and the translation recognition task. The selected words reflected the core target vocabulary of each learning module and were the same for both behavioral and EEG assessments within each class. In one class (Class 2), the instructional units contained fewer than 20 new words per lesson. Therefore, two consecutive learning units were combined, and a subset of 20 items drawn from both units was used for the EEG and behavioral assessments. Separate pre- and post-tests were administered for each learning unit in this class (see also Vocabulary Test).

A subset of 20 items from the vocabulary test was used for the translation recognition task. These items represented the key target words most relevant to the instructional unit, and thus the set of words included in the translation task sometimes differed from that of the vocabulary test (see *Supplementary Table S1* for details).

#### 2.3.2 Vocabulary Test

For each class, 20 target vocabulary items (see Stimuli) were assessed using a handwritten vocabulary test administered in the classroom setting. The tests were completed individually by the children under the supervision of their classroom teacher, both before (pre-test) and after (post-test) the learning intervention.

Children were instructed to provide written translations of each word in the target language. Test sheets were digitized and scored with a custom MATLAB implementation of the Levenshtein edit distance (Levenshtein, 1966; Wagner & Fischer, 1974; The MathWorks, 2022). After lower-casing responses to avoid penalizing capitalization, any response with a distance ≤ 2 from the key was marked correct, permitting minor spelling or typographical deviations while preserving an objective and consistent scoring procedure.

As in the previous study, additional cognitive tests were conducted as part of the overall project protocol (see Skieresz et al., 2025), but these data are not analyzed or discussed here.

#### 2.3.3 Translation Recognition Task

The translation recognition task was adapted from the protocol used in our previous adult study (Skieresz et al., 2025) and based on Pu et al. (2016). On each trial, two words were presented sequentially in black on a light gray background at the center of a 24-inch LCD monitor. Each word appeared for 800 ms, with no interstimulus interval. The intertrial interval was fixed at 1,000 ms and included a jitter of 300–600 ms. Children indicated via keyboard whether the second word was a correct or incorrect translation of the first word. After each response, participants indicated by keypress whether their decision was based on knowledge or guessing.

For each class, all 20 target vocabulary items were presented in both congruent (correct translation) and incongruent (incorrect translation) pairs. Each item appeared once in a congruent and once in an incongruent pairing per block. The task consisted of four blocks, with the direction of translation (L1→L2 and L2→L1) alternated across blocks, and the order of blocks counterbalanced between participants. This resulted in a total of 160 trials per session (20 items × 2 conditions × 4 blocks). Incongruent pairs were fixed across blocks and sessions and were generated by randomly recombining items from the same stimulus set, ensuring semantic mismatch. Task duration ranged from approximately 20 to 30 minutes, with optional short breaks between blocks.

The task was conducted in Octave (version 4.0.3) with PsychToolbox extensions (version 3.0.14) on a Debian system.

### 2.4 Procedure

All assessments were conducted on-site at the respective schools. The vocabulary tests were administered by the classroom teacher in a group setting, shortly before (pre-test) and after (post-test) the classroom-based learning intervention. The duration of the intervention was defined as the interval between the pre- and post-intervention EEG assessments, which closely reflected the length of each instructional unit as scheduled by the classroom teachers. This interval varied between classes (M = 60.5 days, SD = 22.1; see *Supplementary Table S2B* for class-wise values). Cognitive tests (not reported in the present manuscript) were conducted in groups under the supervision of the experimenter, using either school computers or Lenovo laptops provided by UniDistance Suisse.

EEG assessments were performed individually in a separate room within the school building to minimize distractions and ensure standardized conditions. At the pre-intervention EEG session, each child first completed a brief demographic questionnaire and a visual acuity screening using Landolt rings (optikschweiz.ch), followed by electrode setup. Participants were seated approximately 60 cm from the computer screen, and a chin rest was used to minimize head movements and maintain constant viewing distance. Both the pre- and post-intervention EEG sessions included a 3-minute resting-state recording (eyes open), a visual oddball paradigm (with written words presented on the screen), and the translation recognition task, with the order of EEG tasks counterbalanced across participants. Short breaks were provided between tasks as needed, during which the quality of the EEG signal was monitored and adjusted if necessary.

The temporal interval between each classroom-based vocabulary test and the corresponding EEG session (both pre- and post-intervention) also varied across classes, primarily due to differences in curriculum scheduling and logistical constraints at the schools. On average, the pre-intervention EEG was conducted 20.8 days (SD = 3.7) before the pre-vocabulary test, and the post-intervention EEG was conducted 16.3 days (SD = 6.6) before the post-vocabulary test. Class-wise means and standard deviations for these intervals are presented in *Supplementary Table S2B*.

In one class, which covered two consecutive instructional units to reach a minimum of 20 vocabulary items, the test schedule was adjusted accordingly (see Materials).

### 2.5 Electrophysiological Recording and Pre-processing

EEG was recorded using a 64-channel sponge-based electrode system (RNet; Brain Products GmbH), arranged according to the extended International 10–20 system (ground: FPz, reference: Cz). Electrode impedances were kept below 30 kΩ. Signals were digitized at 1,000 Hz using a BrainAmp amplifier (Brain Products GmbH) and recorded on a Lenovo laptop (Windows 10 Home, 8 GB RAM).

Pre-processing was performed in MATLAB using EEGLAB (version 2024.1; Delorme & Makeig, 2004; The MathWorks, Inc., 2022) and ERPLAB (version 12.00; Lopez-Calderon & Luck, 2014). Data were downsampled to 250 Hz and band-pass filtered (0.1–35 Hz). Bad channels were interpolated based on visual inspection. Data were then re-referenced to the common average. To ensure high data quality and maximize trial retention, an adaptive artifact rejection approach was implemented at both the continuous and segmented data level. Specifically, an initial amplitude threshold of ±150 µV was applied, with the threshold increased stepwise (by 25 µV increments, up to a maximum of 400 µV) if more than 20% (continuous) or 50% (epoch-wise) of trials were rejected. This iterative process continued until the proportion of rejected trials fell below the respective cut-off. Preprocessed data were segmented into peri-stimulus epochs (−200 to 800 ms, with −200 to 0 ms baseline), and additional artifact rejection was performed using both amplitude-based and eye-blink detection methods (ERPLAB). After pre-processing, an average of 97.0 (SD = 21.8) pre-learning trials and 96.8 (SD = 23.7) post-learning trials per participant were retained for further analysis.

### 2.6 Data Analysis

#### 2.6.1 Statistical Analyses & Variables

Statistical analyses were conducted in RStudio (version 2024.12.0) using t-tests and mixed ANOVAs, with the significance threshold set at *p* = .05. Behavioral dependent variables included vocabulary test accuracy (percent correct) and response-bias indices from the translation recognition task (sensitivity *d′*, criterion location *c*, and likelihood ratio ln *β*). Response-bias indices were calculated following Macmillan and Creelman (2005), using the correction procedure by Snodgrass and Corwin (1988) to address extreme values, by adding 0.5 and dividing by 𝑁 + 1, where 𝑁 is the number of trials in the corresponding condition (congruent/incongruent).

The specific computations were:

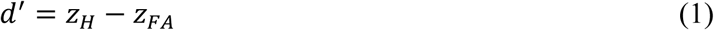

The criterion location was computed as

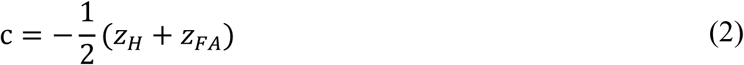

The likelihood ratio was computed as

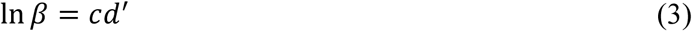

where 𝑧_𝐻_ and 𝑧_𝐹𝐴_ denote the z-transformed hit rate and false alarm rate, respectively.

Electrophysiological dependent variables included mean N400 amplitudes, measured at six centroparietal electrodes (C3, Cz, C4, P3, Pz, P4) in the 300 – 500 ms post-stimulus window, reflecting the canonical centroparietal N400 distribution as established in adults (Kutas & Federmeier, 2011; Van Petten & Luka, 2012). This approach was chosen to ensure consistency with the previous adult study and to enable direct comparison of effect sizes and topographies across age groups, while also aligning with recommendations for ERP analyses in developmental samples (cf. Friedrich & Friederici, 2004). Data were checked for extreme outliers in N400 amplitude and excluded if present (see Results for details).

For subsequent group analyses, participants were classified based on their baseline proficiency, i.e., pre-vocabulary test accuracy, into “low proficiency” and “high proficiency” via a median split within the EEG subsample. For descriptive purposes, participants who completed the vocabulary test but not the EEG sessions (“no EEG data” group) are also reported to illustrate the representativeness of the EEG sample (see *Supplementary Table S2A* for group sizes).

#### 2.6.2 Topographic Clustering, Global Field Power, and Global Map Dissimilarity

Analyses of global features of the EEG were conducted using the freeware Cartool (version 5.02; Brunet et al., 2011) for topographic clustering and the freeware RAGU (compiled 24 November 2020; Koenig et al., 2011) for GFP and GMD.

Topographic clustering decomposes continuous ERPs into transient, but stable, ERP template maps that reflect discrete functional brain states. Critically, group-level template maps derived from the adult sample in our previous study (Skieresz et al., 2025) were directly backfitted to the children’s EEG data and the identical temporal epochs and map configurations were applied (Epoch 1: 0–292 ms; Epoch 2: 292–648 ms; Epoch 3: 648–796 ms), allowing for direct comparison of template-map preponderance across age groups.

Global Field Power (GFP). GFP reflects the overall magnitude of electric field independent of topography and was computed frame-wise (𝑡) across the 0–796 ms as the root-mean-square of each 𝑁 electrode’s deviations from the instantaneous mean across electrodes (Equation 4).

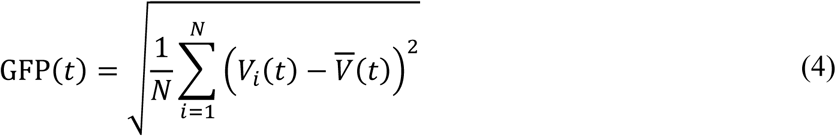

Significance was assessed with cluster-based permutation tests over time (5,000 iterations) and summarized with a global pAUC (time-integrated permutation statistics) over 0 – 796 ms.

Global Map Dissimilarity (GMD/TANOVA). GMD quantified time-resolved topographic differences between conditions after GFP normalization and was computed frame-wise across 0-796 ms. *α*￼*β*￼, GMD *t*￼ was computed as:

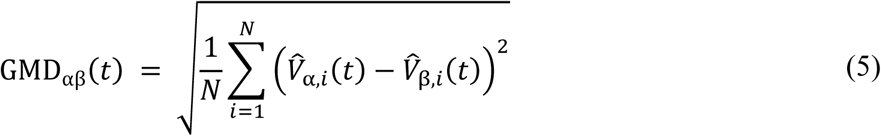

where 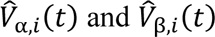 are the GFP-normalized potentials at electrode 𝑖 for conditions 𝛼 and 𝛽, respectively and 𝑁 is the number of electrodes. Time-resolved TANOVA used 5,000-iteration cluster-corrected permutation tests, and we also report the global pAUC over 0 – 796 ms.

For clarity, AUC denotes the time integral of GFP or GMD over 0–796 ms, whereas pAUC in the permutation framework denotes the time-integrated permutation test statistic. All analytical parameters and procedures mirrored those applied in the preceding adult study (Skieresz et al., 2025) to facilitate direct cross-sample comparisons.

#### 2.6.3 Exploratory Associations between Baseline Proficiency and Neural Change

To complement the group analyses, we examined whether baseline proficiency (pre-test vocabulary accuracy, %) correlated with the magnitude of learning-related neural change. For each participant we formed post–pre change scores for the N400 incongruity effect (region of interest; ROI definition as above), the time-integrated global field power (GFP; area under the curve, 0–796 ms), the time-integrated global map dissimilarity (GMD; 0–796 ms), and template-map indices. Template-map indices were summarized as Map 4–Map 3 in Epoch 2 (292–648 ms; N400 window) and Map 6–Map 5 in Epoch 3 (648–796 ms). For these indices, as well as for GFP, values were averaged across congruent/incongruent trials within session before deriving change scores. GMD was computed on GFP-normalized topographies as the time-resolved difference (incongruent–congruent) and then integrated over 0–796 ms per session prior to forming change scores. Associations with baseline proficiency were quantified using two-sided Spearman rank correlations (*ρ*; Spearman, 1904/2010). We report 95% bias-corrected and accelerated (BCa) bootstrap confidence intervals (10,000 resamples; Efron, 1987; Efron & Tibshirani, 1993) and Benjamini–Hochberg false-discovery-rate–adjusted *p* values across the five outcomes (Benjamini & Hochberg, 1995).

## 3 Results

### 3.1 Vocabulary Test

Children’s accuracy in the vocabulary test increased substantially from pre- to post-learning (pre: *M* = 21.0%, *SD* = 18.5, range = 0–75; post: *M* = 61.8%, *SD* = 23.0, range = 10–96.8). This improvement was highly significant, *t*(64) = 14.0, *p* < .001, *d* = 1.70, 95% CI [1.31, 2.07]. For group-level analyses, participants were divided into “Low Proficiency” (N = 20) and “High Proficiency” (N = 22) based on a median split of baseline (pre-vocabulary test) accuracy within the EEG subsample. An additional 23 children completed the vocabulary test, but did not participate in the EEG assessment (“No EEG group”).

**Figure 1.**
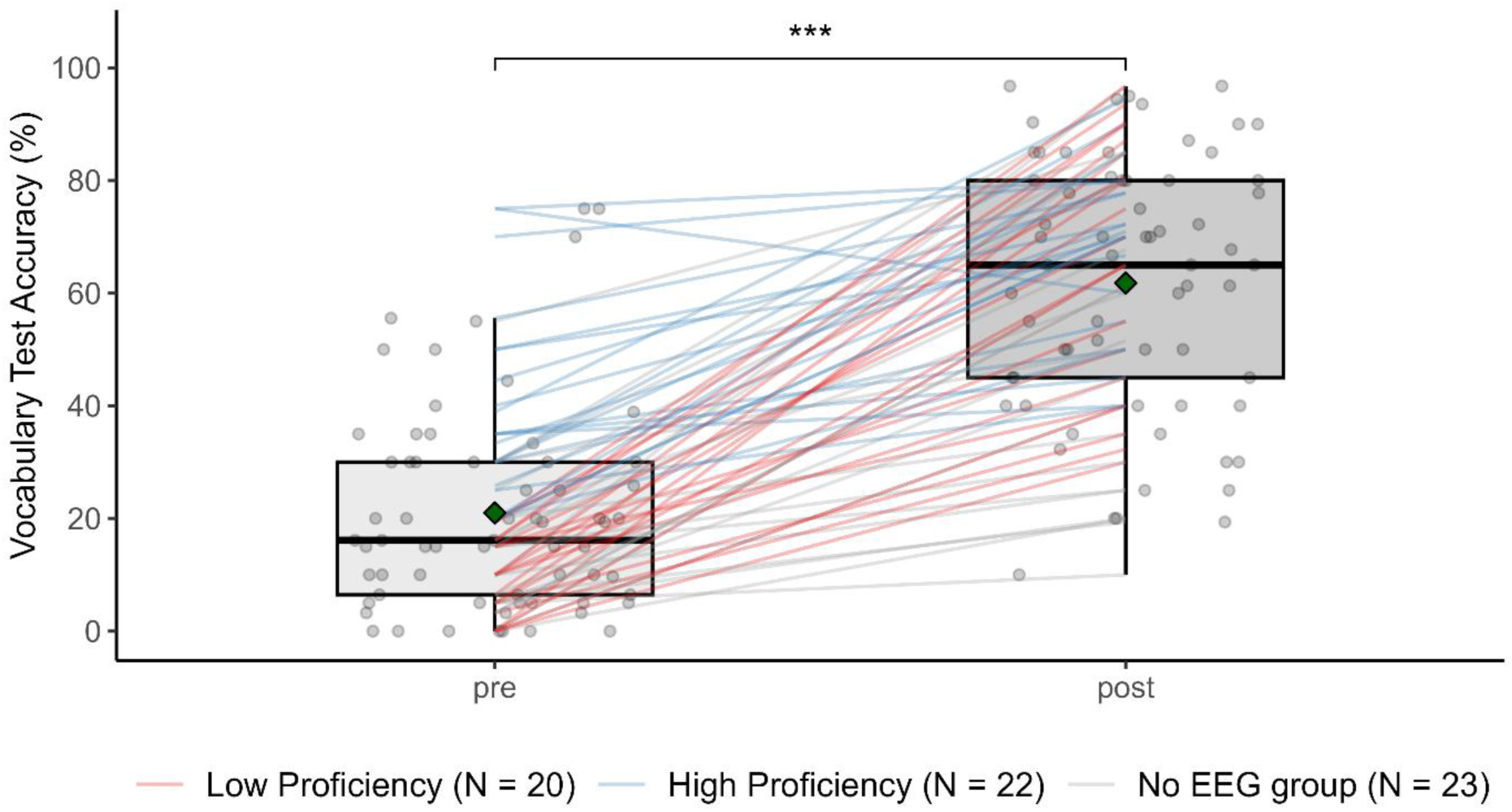
Vocabulary Test Accuracy (Percent Correct) Before and After Learning. *Note.* Boxplots show participants’ test accuracy (%) before and after learning. Gray circles represent individual accuracy. Lines connect individual pre/post measurements, color-coded by proficiency group. Green diamonds indicate group means; asterisks denote significant within-group gains (*p* < .001).

### 3.2 Response-Bias Indices

Sensitivity (*d’*), criterion (*c*) and likelihood ratio (ln *β*) were computed as described in the Methods. Sensitivity in the translation recognition task increased substantially from pre-to post-learning, reflected in a significant main effect of Session, *F*(1, 40) = 92.98, *p* < .001, *η²_g_* = .37, and of Proficiency Group, *F*(1, 40) = 18.51, *p* < .001, *η²_g_* = .26. The low-proficiency group improved from a pre-learning mean of *M* = 1.02 (*SD* = 0.69) to *M* = 2.34 (*SD* = 1.01) after learning, whereas the high-proficiency group increased from *M* = 2.03 (*SD* = 1.08) to *M* = 3.55 (*SD* = 0.99). The interaction was not significant (*p* = .486).

For *c*, there was a significant Session × Proficiency Group interaction, *F*(1, 40) = 7.93, *p* = .008, *η²_g_* = .08. Children with high proficiency showed a reduction in response conservatism after learning (pre: *M* = 0.27, *SD* = 0.33; post: *M* = 0.08, *SD* = 0.27), whereas children with low proficiency became slightly more conservative (pre: *M* = 0.04, *SD* = 0.25; post: *M* = 0.17, *SD* = 0.26).

For ln *β*, the low-proficiency group increased from *M* = 0.08 (*SD* = 0.35) to *M* = 0.39 (*SD* = 0.66), and high-proficiency group decreased from *M* = 0.55 (*SD* = 0.98) to *M* = 0.29 (*SD* = 0.89). No significant main effects or interactions were observed (all *p* ≥ .077).

### 3.3 N400 Incongruity Effect

The N400 incongruity effect was quantified as the amplitude difference between incongruent and congruent trials in the 300–500 ms post-stimulus window at six centroparietal electrodes. We screened ROI amplitudes within each session using a robust median/MAD criterion (absolute robust z ≥ 3.5). One participant with low proficiency exceeded this threshold in the post session (12.13 μV) and was excluded from all analyses (both sessions removed to preserve pairing). A mixed ANOVA with Session (pre, post) and Proficiency Group (low, high) revealed no significant main effect of Session, *F*(1, 39) = 0.93, *p* = .342, *η²_g_* = .01, no significant main effect of Proficiency Group, *F*(1, 39) = 0.31, *p* = .584, *η²_g_* = .00, and no significant Session × Proficiency Group interaction, *F*(1, 39) = 1.09, *p* = .302, *η²_g_* = .01. Thus, there was no evidence for a learning-related increase in the N400 incongruity effect on this ROI analysis.

**Figure 2.**
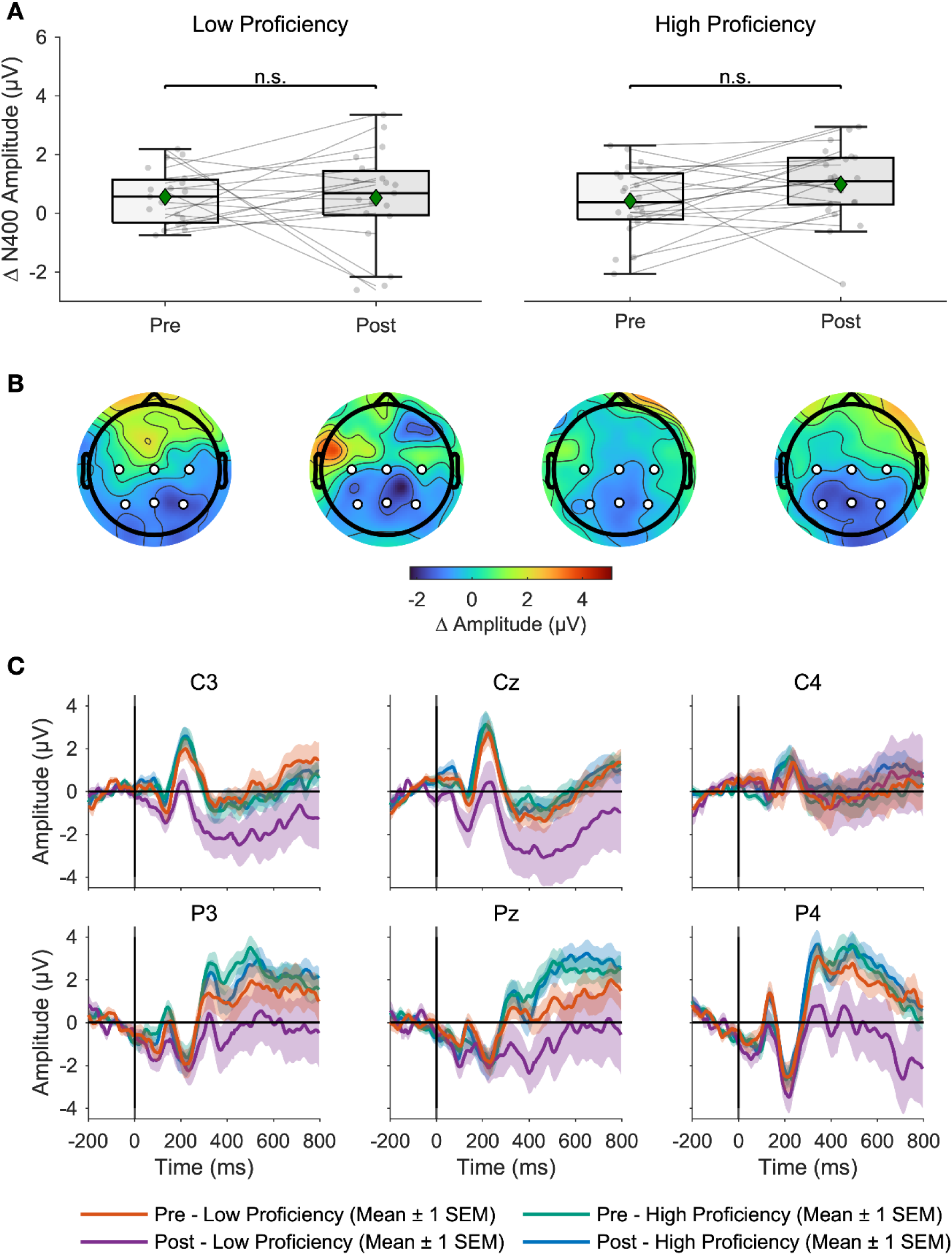
N400 Amplitudes, Topographies, and Grand-Averaged ERPs by Session and Proficiency. *Note.* (A) Mean N400 incongruity amplitudes pre- and post-learning for low (N = 19) and high (N = 22) proficiency. (B) Corresponding scalp topographies with ROI. (C) Grand-averaged ERPs of ROI channels by session and proficiency group (mean ± 1 SEM).

### 3.4 Global Features of the Scalp Electric Field

#### 3.4.1 ERP Template-Map Preponderance

To examine learning-related changes in the strength and topography of the scalp electric field, we backfitted the six adult normative ERP template maps using the same temporal segmentation and epoch definitions (Epoch 1: 0–292 ms, Epoch 2: 292–648 ms, Epoch 3: 648–796 ms). We quantified template-map preponderance as the proportion of time frames assigned to each template map within an epoch.

For the first epoch (0–292 ms; Maps 1 and 2), a mixed ANOVA on template-map preponderance revealed a robust main effect of Map, *F*(1, 39) = 33.77, *p* < .001, *η²_g_* = .30, indicating that Map 2 was generally more dominant than Map 1 across all conditions. No other effects nor interactions reached significance (all *p* ≥ .091).

For the second epoch (292–648 ms; Maps 3 and 4, N400 window), a main effect of Map was observed, *F*(1, 39) = 41.33, *p* < .001, *η²_g_* = .38, indicating greater preponderance of Map 4 than Map 3. In addition, there was a significant interaction between Map and Proficiency Group, *F*(1, 39) = 13.63, *p* = .001, *η²_g_* = .17, and between Congruency and Map, *F*(1, 39) = 5.39, *p* = .026, *η²_g_* = .01. There were no significant main effects of Proficiency Group, Session, or Congruency alone, nor higher-level interactions (all *p* > .09).

To disentangle these effects, follow-up ANOVAs were performed separately for each session and congruency condition. Before learning, both congruent and incongruent trials showed strong main effects of Map (congruent: *F*(1, 39) = 30.59, *p* < .001, *η²_g_* = .44; incongruent: *F*(1, 39) = 15.18, *p* < .001, *η²_g_* = .28), but the Map × Proficiency Group interactions did not reach significance (congruent: *F*(1, 39) = 2.74, *p* = .106; incongruent: *F*(1, 39) = 4.73, *p* = .036, *η²_g_* = .11), with only a modest interaction in the incongruent case.

After learning, both congruent and incongruent conditions revealed significant Map × Proficiency Group interactions. In post-learning congruent trials, there was a main effect of Map (*F*(1, 39) = 31.37, *p* < .001, *η²_g_* = .45) and a significant Map × Proficiency Group interaction (*F*(1, 39) = 9.30, *p* = .004, *η²_g_* = .19), such that the high-proficiency group showed greater Map 4 preponderance and the low-proficiency group greater Map 3 preponderance. The effect was strongest for post-learning incongruent trials: Here, the Map × Proficiency Group interaction reached its largest effect size (*F*(1, 39) = 17.48, *p* < .001, *η²_g_* = .31), alongside a main effect of Map (*F*(1, 39) = 19.01, *p* < .001, *η²_g_* = .33). Thus, the preponderance of Maps 3 and 4 in the N400 time window became strongly proficiency-dependent only after learning, with high-proficiency children showing a shift toward the adult-like map configuration – i.e., greater Map 4 than Map 3 preponderance within Epoch 2 as in our adult reference sample (Skieresz et al., 2025).

In the third epoch (648–796 ms; Maps 5 and 6), a significant three-way interaction was observed between Proficiency Group, Congruency, and Map (*F*(1, 39) = 6.39, *p* = .016, *η²_g_* = .03). Follow-up analyses indicated that this effect was driven specifically by post-learning incongruent trials: High-proficiency group showed greater Map 6 preponderance, whereas low-proficiency group showed greater Map 5 preponderance (*F*(1, 39) = 6.50, *p* = .015, *η²_g_* = .14). No other condition in this epoch showed significant differences (all *p* > .05; see *Supplementary Table S3*).

The preponderance of template maps in Epochs 2 and 3 for all experimental conditions and proficiency groups is visualized in *Figure 3*.

**Figure 3.**
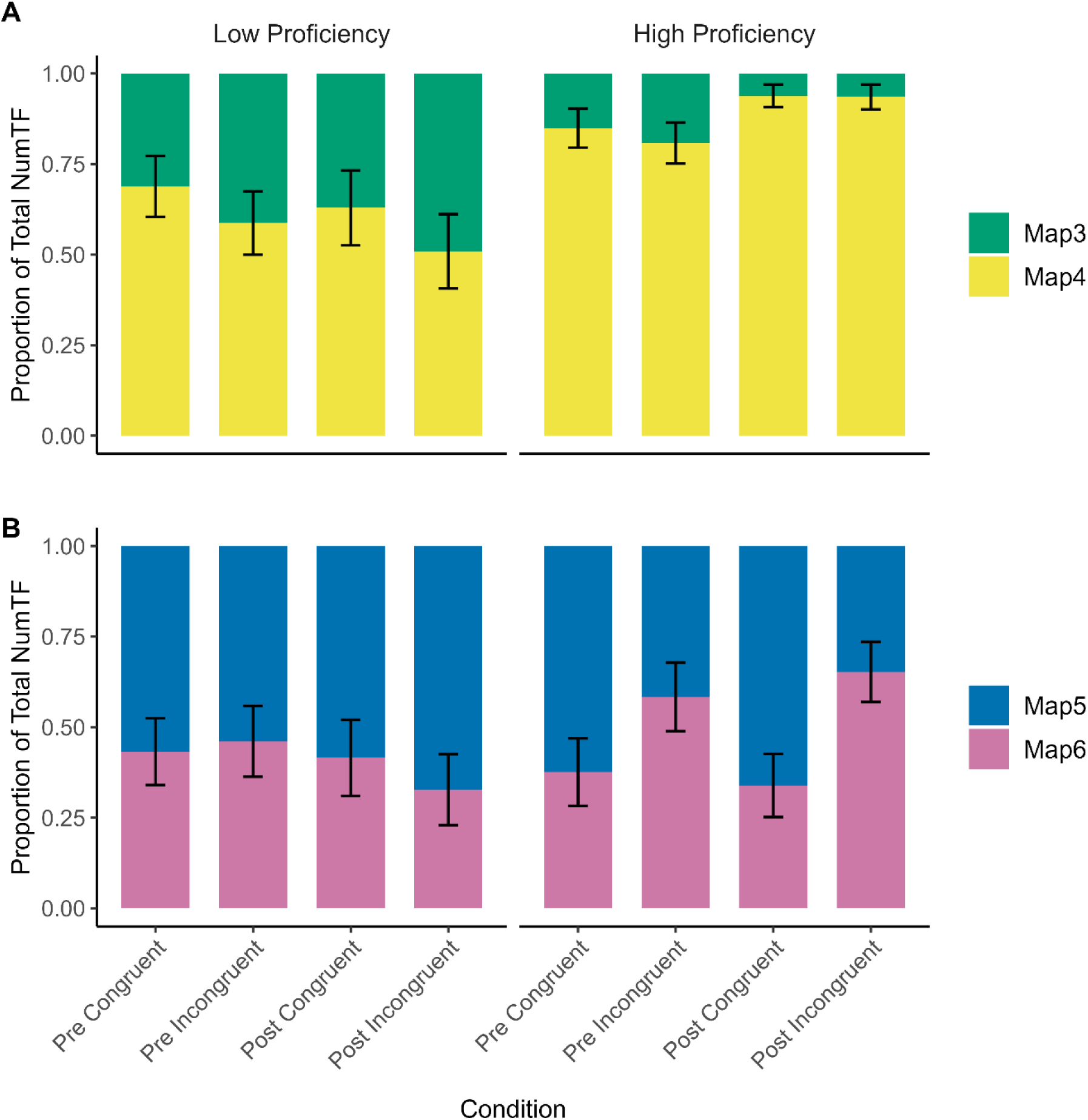
Template-Map Preponderance by Condition and Proficiency Group. *Note.* Template-map preponderance for Map 3 and 4 in Epoch 2 (A), and Map 5 and 6 in Epoch 3 (B) expressed as the proportion of total time frames per condition and proficiency group. Bars represent mean proportions. Error bars indicate the standard error of the mean.

#### 3.4.2 Global Field Power and Global Map Dissimilarity

To complement the ERP template-map analyses (topographic clustering), we examined time-resolved Global Field Power (GFP; response strength) and Global Map Dissimilarity (GMD/TANOVA; topographic change) over the post-stimulus period (0-796 ms).

GFP showed a robust main effect of Session, *p* < .001, and a significant Session × Proficiency Group interaction, *p* < .001, while the main effect of Proficiency Group did not reach conventional significance (*p* = .057; see left column of *Supplementary Figure S1*).

GMD showed main effects for Session (*p* = .047), Proficiency Group (*p* = .005), and their interaction (*p* = .003; see right column of *Supplementary Figure S1*).

These results indicate proficiency-dependent reorganization of scalp topographies with learning. These effects extended beyond the N400 window. Cluster-corrected time courses are visualized in *Supplementary Figure S1*.

### 3.5 Baseline proficiency as a Correlate of Neural Change

In an exploratory correlational analysis, we tested whether baseline proficiency (pre-vocabulary test accuracy) correlated with the magnitude of learning-related neural change. Across outcomes – ΔGFP AUC, ΔGMD AUC, ΔN400, Δ(Map 4 − Map 3), and Δ(Map 6 − Map 5) – correlations were small (|*ρ*| ≤ .25) and non-significant after FDR correction; the largest was for ΔGFP AUC (*ρ* = −.25, 95% BCa CI [−.51, .07], *p* = .116, *p*FDR = .580). Full statistics are provided in *Supplementary Table S4*.

## 4 Discussion

Children’s classroom-based L2 vocabulary learning was examined to determine whether an adult normative set of ERP template maps would characterize schoolchildren’s ERPs and to assess learning-related change using both a canonical centroparietal N400 incongruity measure and electrical-neuroimaging indices. We recorded 64-channel EEG during a translation-recognition task before and after a regular instructional unit, grouped children by baseline proficiency, and quantified the N400 (300–500 ms, centroparietal ROI) alongside template-map preponderance, Global Field Power (GFP), and Global Map Dissimilarity (GMD). Template-map preponderance was derived by backfitting the adult template set. The adult templates backfitted cleanly across sessions and conditions, indicating a shared canonical topographic configuration across age groups. The N400 incongruity effect did not increase from pre- to post-learning, whereas topographic and global measures showed robust session effects and proficiency dependencies: within the N400 window, template-map preponderance diverged by proficiency (higher Map 4, lower Map 3 in higher-proficiency children), and a later interval (∼650–800 ms) also differed by proficiency and congruency. Thus, in this naturalistic context, learning-related neural changes were more evident in scalp topography and global field dynamics than in a single ROI-based N400 amplitude, motivating the subsequent comparison to our adult reference sample.

### 4.1 Relating the Findings to an Adult Reference Sample

In our child sample, the grand-average ERPs show a clearly time-locked N400 in the canonical 300–500 ms window without evidence for a delayed peak or a systematic anterior shift (*Figure 2C*). What distinguishes the children – particularly in the post-learning low-proficiency group – is the more diffuse scalp pattern and broader variability (large SEM bands), rather than latency differences (at least as characterized here). This profile is compatible with classic developmental work showing that school-aged children often exhibit broader and sometimes slightly anterior N400 distributions, even when their timing is broadly adult-like (Coch & Holcomb, 2003; Friedrich & Friederici, 2004; Holcomb et al., 1992; Juottonen et al., 1996; see also Kutas & Federmeier, 2011). At the same time, N400 topography varies with task difficulty and proficiency: more demanding or less familiar conditions tend to yield more fronto-central and diffuse patterns, whereas easier or better-learned items show the canonical posterior distribution (Coch & Holcomb, 2003; Elgort et al., 2014; Kutas & Federmeier, 2011; Perrin & García-Larrea, 2003; Van Petten & Luka, 2012; Yum et al., 2014).

In our data, the diffuse scalp pattern and larger variability – especially in post-learning low-proficiency group– thus likely index residual difficulty rather than a latency shift. Against this backdrop, our learning effects manifested less as a univariate amplitude increase and more as changes in global strength and in the preponderance of template maps (Murray et al., 2008). In Epoch 2 (the N400 window), Map 4 predominated overall, but its preponderance depended on both proficiency and congruency after learning – high-proficiency children showed a greater preponderance of Map 4, whereas low-proficiency children showed greater preponderance of Map 3 – a divergence that was not present pre-learning. Relative to our adult reference sample (Skieresz et al., 2025), the children showed a weaker congruency-based separation of maps and a stronger dependence on individual proficiency; similarly, late effects in Epoch 3 were clearest for post-learning incongruent items and were most consistent in high-proficiency group, whereas adults showed a more uniform late-epoch pattern (Map 6 for incongruent and a tendency toward Map 5 for congruent), even though the latter was not always statistically significant.

This combination – adult-like N400 timing with more diffuse topographies and proficiency-modulated template-map preponderance – fits broader developmental evidence that children’s resting-state EEG template maps tend to be shorter, less stable, and more variable than adults’ (Koenig et al., 2002; Tomescu et al., 2018), which likely attenuates the condition-wise map separation seen in mature systems.

### 4.2 Topographic Dynamics and Global Measures

Topographic ERP template-map analyses provide a window into transient, quasi-stable functional states that tile the ERP time course. In the present data, the N400 window was dominated by two maps whose preponderance flipped as a function of proficiency after learning (cf. Brandeis et al., 1995, who reported distinct pre-N400 and N400 scalp maps during sentence processing). This aligns with developmental work showing that the basic topographic classes are present in childhood, but states are shorter, less stable, and transition more frequently than in adults, with occupancy and stability increasing through adolescence (Koenig et al., 2002; Tomescu et al., 2018). Within this framework, our findings suggest that learning pushes high-proficiency children into an adult-like map configuration (greater Map 4 preponderance in Epoch 2), whereas low-proficiency children spend comparatively more time in a pre-learning configuration – even when overall task performance improves. The GFP modulation across sessions, together with its interaction with proficiency, is consistent with either larger underlying source activity and/or improved temporal precision (i.e., reduced trial-to-trial latency variability) after learning; GFP alone cannot disambiguate these mechanisms (see Michel & Koenig, 2018; Murray et al., 2008). Critically, concurrent GMD effects—which are insensitive to overall field strength—indicate a proficiency-dependent reorganization of the underlying generator configuration, not merely a uniform gain change. These observations dovetail with the view of the N400 as a family of distributed processes supporting semantic access and integration (Kutas & Federmeier, 2011; Van Petten & Luka, 2012) and with electrical-neuroimaging demonstrations that learning can modify both response strength and the recruitment of distinct networks (Murray et al., 2008).

### 4.3 Individual Differences as a Driver of Neural Specialization

Proficiency groups were defined via a median split on baseline vocabulary accuracy; accordingly, the high-proficiency group knew more words at pre-test. At the neural level, they also showed larger session-related increases in GFP and a shift in ERP template-map preponderance toward an adult-like configuration within the N400 window, together with a characteristic late-epoch pattern in incongruent trials. We therefore interpret these group differences as Session × Proficiency effects in neural measures rather than evidence that higher baseline accuracy linearly predicts greater neural change across individuals. Consistent with this interpretation, exploratory correlations between baseline proficiency and neural change scores (ΔN400, ΔGFP AUC, ΔGMD AUC, Δ[Map 4–Map 3], Δ[Map 6–Map 5]) were small and non-significant after FDR correction (|*ρ*| ≤ .25; all *p*FDR ≥ .58; see Section 3.5 and *Supplementary Table S4*).

This proficiency-contingent pattern is consistent with greater efficiency and reliability of semantic access as knowledge consolidates – i.e., faster, more stable engagement of the appropriate networks – aligning with proficiency effects in the adult literature (Elgort et al., 2014; Kutas & Federmeier, 2011; Van Petten & Luka, 2012) and with accounts linking N400 dynamics to knowledge-dependent prediction error (Rabovsky et al., 2018). It also aligns with developmental evidence that children with higher language competence tend to exhibit more focal, adult-like N400 topographies, whereas lower-proficiency peers show broader, more variable negativities (Benau et al., 2011; Coch & Holcomb, 2003; Hahne et al., 2004; Weber-Fox & Neville, 1996).

From a topographic-mapping perspective, higher preponderance of the adult-like template among high-proficiency children suggests more stable recruitment of task-relevant large-scale networks, in line with developmental evidence for increasingly focal and stable scalp topographies with age (Koenig et al., 2002; Tomescu et al., 2018).

Although we did not analyze translation direction separately, the pattern is compatible with the Revised Hierarchical Model, whereby growing L2 knowledge strengthens semantic mediation during translation recognition (Kroll & Stewart, 1994).

### 4.4 Why Did an N400 Incongruity Effect Not Emerge

It is tempting to treat the N400 incongruity effect as the definitive neural signature of successful L2 word acquisition, given extensive evidence for its reliability in adult laboratory work and translation paradigms (Kutas & Federmeier, 2011; Midgley et al., 2009; Pu et al., 2016; Skieresz et al., 2025).

Several features of the present developmental, classroom-based context argue against that expectation. First, children’s N400 scalp distributions are typically broader (and, in some reports, slightly more anterior) than in adults’ (Friedrich & Friederici, 2004; Holcomb et al., 1992; Juottonen et al., 1996; see also Kutas & Federmeier, 2011), although in the present sample the distribution was diffuse rather than anterior (*Figure 2B*). Accordingly, a fixed centroparietal ROI can under-sample the effect when children’s topographies are broad and variable across individuals.

Second, developmental differences in state properties – shorter, less stable, and more variable topographies (Koenig et al., 2002; Tomescu et al., 2018) – tend to dilute condition differences when averaging amplitudes over a fixed 300–500 ms window.

Third, the ecological constraints that make this study informative also introduce heterogeneity that can dampen amplitude sensitivity: the EEG task drew on class-specific subsets of the written vocabulary test, typographical outliers were excluded on a task-specific basis, and the lag between classroom testing and EEG varied across classes.

None of these factors negates the presence of learning—demonstrated behaviorally and by robust session and session-by-proficiency effects in GFP and GMD—but they can attenuate a single amplitude contrast when it is summarized over a fixed six-electrode centroparietal ROI.

The implication is methodological rather than negative: in developmental and ecologically rich settings, electrical-neuroimaging measures that track where and for how long specific topographies prevail, alongside global field strength and topographic dissimilarity, provide a more sensitive assay of learning-related change than a canonical N400 amplitude taken from a fixed six-electrode centroparietal ROI. (Michel & Koenig, 2018; Murray et al., 2008; Van Petten & Luka, 2012).

### 4.5 Limitations

This study has several limitations that qualify the interpretation and generalizability of the findings. First, we backfitted adult normative ERP template maps to children’s data to maximize comparability with our prior adult study (Skieresz et al., 2025). The fact that these templates backfitted well in children supports a set of shared canonical scalp configurations across age groups; nevertheless, future work should also derive child-specific template sets and directly contrast them with adult solutions to test for developmental specializations that an imposed adult solution may obscure.

Second, materials and measurement logistics reflected authentic classroom constraints. The vocabulary items used in the handwritten tests, and the EEG task were only partially harmonized, the number of analyzable trials was limited by having ∼20 target items, and delays between classroom testing and EEG sessions varied across classes. Such heterogeneity – together with natural fluctuations in attention and motivation – can dilute amplitude-based contrasts (e.g., a fixed centroparietal N400 ROI) and inflate between-participant variance, even as global/topographic metrics remain informative.

Third, several methodological choices may have shaped sensitivity. We used a fixed centroparietal ROI and a 300–500 ms window defined a priori from adult studies; broader or slightly shifted child topographies can be undersampled by this choice. Proficiency groups were defined by a median split, which is practical but coarser than continuous modeling. EEG was acquired with a sponge-based 64-channel system and adaptive artifact thresholds (up to ±400 µV) to preserve trials in a school setting; while pragmatic, these steps may introduce residual noise heterogeneity across participants. Moreover, the exploratory proficiency-change analyses were conservative and likely underpowered (EEG N = 41); the small number of items on the vocabulary test, class-specific item sets, and nonzero lags between behavioral and EEG sessions may have attenuated monotonic associations with change scores.

Beyond these limitations, the present study links neurophysiological dynamics to observable learning outcomes in an authentic classroom setting. In doing so, it refines and empirically tests developmental accounts of language processing – including neurocognitive models of semantic integration (e.g., Friederici, 2006; Kutas & Federmeier, 2011) – by mapping age- and proficiency-related differences in electrophysiological indices of L2 acquisition. These considerations point toward age-appropriate, evidence-based assessment and intervention and support translational research at the interface of neuroscience and education.

### 4.6 Conclusion

In school-aged children learning L2 vocabulary in real classrooms, learning-related neural change was clearer in scalp topography and global field dynamics than in a single centroparietal N400 incongruity amplitude. Learning increased GFP and redistributed template-map preponderance in a proficiency-dependent fashion within – and beyond – the canonical N400 window. These results align with developmental evidence on the maturation of ERP template maps and N400 topographies and support an analysis strategy that treats the N400 not merely as an amplitude at a few electrodes, but as a distributed, temporally structured family of states whose prevalence (preponderance) and strength track both learning and proficiency (Koenig et al., 2002; Kutas & Federmeier, 2011; Michel & Koenig, 2018; Murray et al., 2008).

Practically, electrical neuroimaging offers sensitive, ecologically valid biomarkers of vocabulary learning in children, complementing – and in some contexts surpassing – canonical ERP amplitudes.

## Author Contributions

Nicole H. Skieresz: Conceptualization; Methodology; Investigation; Formal analysis; Data curation; Visualization; Writing – original draft; Writing – review & editing.

Sandy C. Marca: Investigation; Data curation.

Micah M. Murray: Supervision; Conceptualization; Methodology; Writing – review & editing; Funding acquisition.

Nicolas Rothen: Supervision; Conceptualization; Methodology; Writing – review & editing; Funding acquisition; Project administration.

Thomas P. Reber: Supervision; Conceptualization; Methodology; Writing – review & editing; Funding acquisition; Project administration.

*Note.* Micah M. Murray, Nicolas Rothen, and Thomas P. Reber contributed equally to this work (*).

## Data Availability Statement

All data and analysis scripts are available on the Open Science Framework (OSF): https://osf.io/e3jkc/?view_only=40f2f0ba7b6041c6b0b6a1be4df21532.

## References

Ansari, D., & Coch, D. (2006). Bridges over troubled waters: Education and cognitive neuroscience. Trends in Cognitive Sciences, 10(4), 146–151. 10.1016/j.tics.2006.02.007

Benau, E. M., Morris, J., & Couperus, J. W. (2011). Semantic Processing in Children and Adults: Incongruity and the N400. Journal of Psycholinguistic Research, 40, 225–239. 10.1007/s10936-011-9167-1

Benjamini, Y., & Hochberg, Y. (1995). Controlling the false discovery rate: A practical and powerful approach to multiple testing. Journal of the Royal Statistical Society: Series B (Methodological), 57(1), 289–300. 10.1111/j.2517-6161.1995.tb02031.x

Brandeis, D., Lehmann, D., Michel, C. M., & Mingrone, W. (1995). Mapping event-related brain potential microstates to sentence endings. Brain Topography, 8(2), 145–159. 10.1007/BF01199778

Brunet, D., Murray, M. M., & Michel, C. M. (2011). Cartool: A software for the visualization and analysis of multichannel EEG data. Retrieved from https://cartool.cibm.ch

Coch, D., & Holcomb, P. J. (2003). The N400 in beginning readers. Developmental Psychobiology, 43(2), 146–166. 10.1002/dev.10129

Delorme, A., & Makeig, S. (2004). EEGLAB: An open-source toolbox for analysis of single-trial EEG dynamics. Journal of Neuroscience Methods, 134(1), 9–21. 10.1016/j.jneumeth.2003.10.009

Dikker, S., Wan, L., Davidesco, I., Kaggen, L., Oostrik, M., McClintock, J., Rowland, J., Michalareas, G., Van Bavel, J. J., Ding, M., Poeppel, D., & Hasson, U. (2017). Brain-to-brain synchrony tracks real-world dynamic group interactions in the classroom. Current Biology, 27(9), 1375–1380. 10.1016/j.cub.2017.04.002

Efron, B. (1987). Better bootstrap confidence intervals. Journal of the American Statistical Association, 82(397), 171–185. 10.1080/01621459.1987.10478410

Efron, B., & Tibshirani, R. J. (1993). An introduction to the bootstrap. Chapman & Hall/CRC. 10.1201/9780429246593

Elgort, I., Perfetti, C. A., Rickles, B., & Stafura, J. Z. (2014). Contextual learning of L2 word meanings: second language proficiency modulates behavioural and event-related brain potential (ERP) indicators of learning. Language, Cognition and Neuroscience, 30(5), 506–528. 10.1080/23273798.2014.942673

Friederici, A. D. (2006). The neural basis of language development and its impairment. Neuron, 52(6), 941–952. 10.1016/j.neuron.2006.12.002

Friedrich, M., & Friederici, A. D. (2004). N400-like Semantic Incongruity Effect in 19-Month-Olds: Processing Known Words in Picture Contexts. Journal of Cognitive Neuroscience, 16(8), 1465–1477. 10.1162/0898929042304705

Hahne, A., Eckstein, K., & Friederici, A. D. (2004). Brain signatures of syntactic and semantic processes during children’s language development. Journal of cognitive neuroscience, 16(7), 1302–1318. 10.1162/0898929041920504

Holcomb, P. J., Coffey, S. A., & Neville, H. J. (1992). Visual and auditory sentence processing: A developmental analysis using event-related brain potentials. Developmental Neuropsychology, 8(2-3), 203–241. 10.1080/87565649209540525

Juottonen, K., Revonsuo, A., & Lang, H. (1996). Dissimilar age influences on two ERP waveforms (LPC and N400) reflecting semantic context effect. Brain research. Cognitive brain research, 4(2), 99–107. 10.1016/0926-6410(96)00022-5

Koenig, T., Kottlow, M., Stein, M., & Melie-García, L. (2011). RAGU: A free tool for the analysis of EEG/MEG data using randomisation statistics. Brain Topography, 24(1), 3–12. 10.1007/s10548-010-0177-1

Koenig, T., Lehmann, D., Michel, C. M., & Brandeis, D. (2002). Millisecond by millisecond, year by year: Normative EEG microstates and developmental stages. NeuroImage, 16(1), 41–48. 10.1006/nimg.2002.1070

Kroll, J. F., & Stewart, E. (1994). Category interference in translation and picture naming: Evidence for asymmetric connections between bilingual memory representations. Journal of Memory and Language, 33(2), 149–174. 10.1006/jmla.1994.1008

Kutas, M., & Federmeier, K. D. (2011). Thirty years and counting: Finding meaning in the N400 component of the eventrelated brain potential (ERP). Annual Review of Psychology, 62, 621–647. 10.1146/annurev.psych.093008.131123

Levenshtein, V. I. (1966). Binary codes capable of correcting deletions, insertions, and reversals. Soviet Physics Doklady, 10(8), 707–710.

Lopez-Calderon, J., & Luck, S. J. (2014). ERPLAB: An open-source toolbox for the analysis of event-related potentials. Frontiers in Human Neuroscience, 8, 213. 10.3389/fnhum.2014.00213

Macmillan, N. A., & Creelman, C. D. (2005). Detection theory: A user’s guide (2nd ed.). Lawrence Erlbaum Associates Publishers.

Michel, C. M., & Koenig, T. (2018). EEG microstates as a tool for studying the temporal dynamics of whole-brain neuronal networks: A review. NeuroImage, 180(Pt B), 577–593. 10.1016/j.neuroimage.2017.11.062

Michel, C. M., Seeck, M., & Murray, M. M. (2004). The speed of visual cognition. Supplements to Clinical Neurophysiology, 57, 617–627. 10.1016/S1567-424X(09)70401-5

Midgley, K. J., Holcomb, P. J., & Grainger, J. (2009). Masked repetition and translation priming in second language learners: A window on the time-course of form and meaning activation using ERPs. Psychophysiology, 46(3), 551–565. 10.1111/j.1469-8986.2009.00784.x

Murray, M. M., Brunet, D., & Michel, C. M. (2008). Topographic ERP analyses: a step-by-step tutorial review. Brain topography, 20(4), 249–264. 10.1007/s10548-008-0054-5

Neville, H. J., Coffey, S. A., Holcomb, P. J., & Tallal, P. (1993). The neurobiology of sensory and language processing in language-impaired children. Journal of cognitive neuroscience, 5(2), 235–253. 10.1162/jocn.1993.5.2.235

Perrin, F., & García-Larrea, L. (2003). Modulation of the N400 potential during auditory phonological/semantic interaction. Cognitive Brain Research, 17(1), 36–47. 10.1016/s0926-6410(03)00078-8

Pu, H., Holcomb, P. J., & Midgley, K. J. (2016). Neural changes underlying early stages of L2 vocabulary acquisition. Journal of Neurolinguistics, 40, 55–65. 10.1016/j.jneuroling.2016.05.002

R Core Team. (2024). R: A language and environment for statistical computing (Version 2024.12.0) [Computer software]. R Foundation for Statistical Computing. https://www.R-project.org/

Rabovsky, M., Hansen, S. S., & McClelland, J. L. (2018). Modelling the N400 brain potential as change in a probabilistic representation of meaning. Nature Human Behaviour, 2(9), 693–705. 10.1038/s41562-018-0406-4

Schlaggar, B. L., Brown, T. T., Lugar, H. M., Visscher, K. M., Miezin, F. M., & Petersen, S. E. (2002). Functional neuroanatomical differences between adults and school-age children in the processing of single words. Science, 296(5572), 1476–1479. 10.1126/science.1069464

Skeide, M. A., Brauer, J., & Friederici, A. D. (2015). Brain functional and structural predictors of language performance. Cerebral Cortex, 26(5), 2127–2139. 10.1093/cercor/bhv042

Skieresz, N. H., Marca, S. C., Murray, M., Reber, T. P., Rothen, N. (2025). Brain Network Differences in Second Language Learning Depend on Individual Competencies. bioRxiv. 10.1101/2025.09.28.679014

Snodgrass, J. G., & Corwin, J. (1988). Pragmatics of measuring recognition memory: Applications to dementia and amnesia. Journal of Experimental Psychology: General, 117(1), 34–50. 10.1037/0096-3445.117.1.34

Spearman, C. (2010). The proof and measurement of association between two things. International Journal of Epidemiology, 39(5), 1137–1150. (Original work published 1904). 10.1093/ije/dyq191

The MathWorks, Inc. (2022). MATLAB (Version R2022b) [Computer software]. The MathWorks, Inc.

Tomescu, M. I., Rihs, T. A., Rochas, V., Hardmeier, M., Britz, J., Allali, G., Fuhr, P., Eliez, S., & Michel, C. M. (2018). From swing to cane: Sex differences of EEG resting-state temporal patterns during maturation and aging. Developmental cognitive neuroscience, 31, 58–66. 10.1016/j.dcn.2018.04.011

Van Petten, C., & Luka, B. J. (2012). Prediction during language comprehension: Benefits, costs, and ERP components. International Journal of Psychophysiology, 83(2), 176–190. 10.1016/j.ijpsycho.2011.09.015

Wagner, R. A., & Fischer, M. J. (1974). The string-to-string correction problem. Journal of the ACM, 21(1), 168–173. 10.1145/321796.321811

Weber-Fox, C., & Neville, H. J. (1996). Maturational constraints on functional specializations for language processing: ERP and behavioral evidence in bilingual speakers. Journal of Cognitive Neuroscience, 8(3), 231–256. 10.1162/jocn.1996.8.3.231

Yum, Y. N., Midgley, K. J., Holcomb, P. J., & Grainger, J. (2014). An ERP study on initial second language vocabulary learning. Psychophysiology, 51(4), 364–373. 10.1111/psyp.12183

