## Supplementary Material for "Proficiency-Dependent and Dynamic Reorganization of Brain Networks During Children’s Second Language Learning"

**Table S1**

*Stimuli by Task and Class*

*Table 1 (vocabulary lists)*

All original vocabulary test items, recognition task subsets, and their English translations are openly available on the Open Science Framework (OSF):

https://osf.io/e3jkc/?view_only=40f2f0ba7b6041c6b0b6a1be4df21532

Readers interested in the exact stimulus materials can download them directly

from this repository or contact the corresponding author.

**Table S2A**

*Class-wise demographic and sample characteristics*

| **Class (grade)** | **L1-L2** | **N** | **Female** | **Age** *M* (*SD*) | **Age Range (min – max)** | **N EEG** | **Female EEG** | **Age EEG** *M* (*SD*) | **Age EEG Range** |
|---|---|---|---|---|---|---|---|---|---|
| Class 1 (4th) | DeFr | 3 | 2 | 10.0 (0.0) | 10 – 10 | 2 | 1 | 10.0 (0.0) | 10 – 10 |
| Class 2 (4th) | DeFr | 13 | 4 | 10.1 (0.4) | 10 – 11 | 7 | 1 | 10.1 (0.4) | 10 – 11 |
| Class 3 (4th) | DeFr | 11 | 6 | 10.0 (0.0) | 10 – 10 | 9 | 6 | 10.0 (0.0) | 10 – 10 |
| Class 4 (4th) | DeFr | 11 | 8 | 10.4 (0.5) | 10 – 11 | 5 | 2 | 10.4 (0.5) | 10 – 11 |
| Class 5 (4th) | FrDe | 5 | 2 | 9.4 (0.5) | 9 – 10 | 5 | 2 | 9.4 (0.5) | 9 – 10 |
| Class 6 (6th) | DeFr | 16 | 8 | 12.0 (0.7) | 11 – 13 | 8 | 3 | 12.0 (0.5) | 11 – 13 |
| Class 7 (6th) | FrDe | 6 | 4 | 11.8 (1.6) | 11 – 15 | 6 | 4 | 11.8 (1.6) | 11 – 15 |

*Note.* Classes are anonymized. DeFr = German native speakers learning French; FrDe = French native speakers learning German; 4th = 4th grade; 6th = 6th grade; "N EEG" = number of children with pre/post EEG data.

**Table S2B**

*Class-wise intervention duration and assessment intervals (in days)*

| Class | EEG Interval (days) *M* (*SD*) | PreVoci-PreEEG interval (days) *M* (*SD*) | PostVoci-PostEEG interval (days) *M* (*SD*) |
|---|---|---|---|
| Class 1 | 49.0 (0) | -21.0 (0) | 1.0 (0) |
| Class 2 | 91.3 (1.9) | -0.1 (2.7) | -4.9 (2.0) |
| Class 3 | 70.0 (15.4) | 12.6 (16.6) | -5.9 (6.7) |
| Class 4 | 44.0 (7.0) | 29.0 (3.0) | -19.0 (9.6) |
| Class 5 | 34.8 (0.4) | -13.6 (0.9) | -23.8 (0.4) |
| Class 6 | 72.1 (5.6) | -21.3 (2.2) | -47.1 (5.0) |
| Class 7 | 34.0 (0.9) | -6.3 (0.5) | -14.7 (22.8) |

*Note.* “EEG interval” = duration between pre- and post-EEG assessment. “PreVoci–PreEEG” = days between pre-vocabulary test and pre-EEG assessment; “PostVoci–PostEEG” = days between post-vocabulary test and post-EEG reassessment. Negative values indicate that the EEG session took place before the respective vocabulary test.

**Table S3**

*Microstate Duration by Map, Session, Congruency, and Proficiency Group*

| Model | Predictor | *F (df1, df2)* | *p* | $\eta^2_g$ |
|---|---|---|---|---|
| **Epoch 1 (TF 1 – 73)** | Map | 33.77 (1, 39) | < .001 | .30 |
| | Proficiency Group × Map | 3.00 (1, 39) | .091 | .04 |
| **Epoch 2 (TF 74 – 163)** | Map | 41.33 (1, 39) | < .001 | .38 |
| | Proficiency Group × Map | 13.63 (1, 39) | .001 | .17 |
| | Congruency × Map | 5.39 (1, 39) | .026 | .01 |
| | Proficiency Group × Session × Map | 2.39 (1, 39) | .095 | .02 |
| **Epoch 3 (TF 164 – 199)** | Congruency × Map | 4.03 (1, 39) | .052 | .02 |
| | Proficiency Group × Congruency × Map | 6.39 (1, 39) | .016 | .03 |

*Note***.** *F*(1, 75) refers to the F-statistic with degrees of freedom (*df1, df2*). *df1* indicates degrees of freedom numerator. *df2* indicates degrees of freedom denominator. *p* values are two-tailed. $\eta^2_g$ = generalized eta squared effect size measure.

**Table S4**

*Associations between Baseline Proficiency and Neural Change (Spearman correlations)*

| Outcome | *ρ* | 95% BCa CI | *p* | *p*FDR |
|---|---|---|---|---|
| ΔN400 ROI (300–500 ms) | 0.16 | [−0.16, 0.47] | .311 | .692 |
| Δ(Map 4 − Map 3) (Epoch 2) | 0.10 | [−0.21, 0.41] | .522 | .692 |
| Δ(Map 6 − Map 5) (Epoch 3) | 0.04 | [−0.30, 0.38] | .816 | .816 |
| ΔGFP (AUC 0–796 ms) | −0.25 | [−0.51, 0.07] | .116 | .580 |
| ΔGMD (AUC 0–796 ms) | −0.10 | [−0.39, 0.22] | .553 | .692 |

*Note.* 95% CIs are BCa from 10,000 bootstrap resamples. *p* values are two-sided; *p*FDR uses the Benjamini–Hochberg procedure across the five outcomes. Δ denotes post–pre change. AUC = time integral (0–796 ms). *ρ* = Spearman's rho.

**Figure S1**

*Significant Effects in GFP and GMD Across the Time Course*

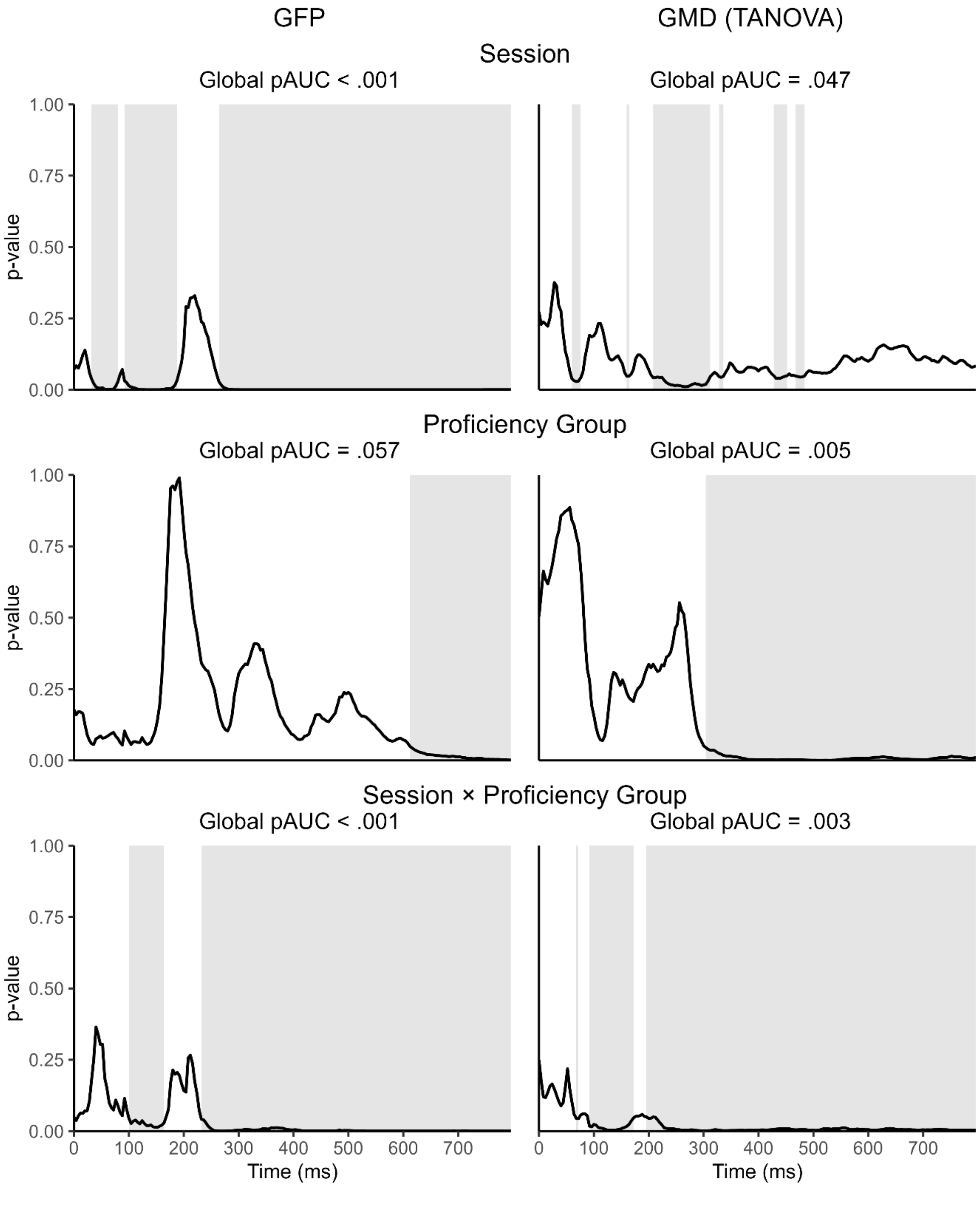


*Note.* Cluster-based permutation p-value time courses with global pAUC in GFP and GMD/TANOVA. Shaded regions indicate significant clusters ($p < .05$). Global pAUC values are shown above each plot.
